# Silk fibroin as a modulator of the amyloidogenic α-synuclein aggregation

**DOI:** 10.1101/2025.10.25.684501

**Authors:** Marco E. Miali, Yoav Barak, Yael Fridmann-Sirkis, Irit Goldian, Ori Brookstein, Dror Eliaz, Ulyana Shimanovich

## Abstract

The aggregation of fiber-forming proteins presents a duality of biological functions, where certain proteins, such as α-synuclein, are implicated in neurodegenerative disorders like Parkinson’s, while others, like silk fibroin, serve as essential building blocks for functional biomaterials. Despite ongoing investigations into the interplay between these contrasting protein groups, existing reports suggest that the presence of “functional” fibrillar proteins can accelerate amyloid growth via a crowding effect. In this study, we report a counterintuitive outcome, demonstrating that silk fibroin inhibits amyloidogenic aggregation of α-synuclein protein. Our findings reveal that although the fibrillar aggregation of both proteins—individually and in mixtures—remains affected by environmental factors such as concentration and temperature, silk fibroin significantly alters the rheological behavior of the mixed solutions. Our structural and kinetic analyses reveal that silk modifies the structural transitions and self-assembly dynamics of α-synuclein through macromolecular crowding and non-specific interactions that eventually suppressed the amyloidogenic aggregation of the α-synuclein. Notably, when nanofibrillar amyloidogenic assemblies are forced to form in the presence of silk, they exhibit a markedly increased susceptibility to enzymatic degradation, a phenomenon not observed with pure α-synuclein fibrillar constructs. These results prompt further investigation into the potential role of “functional” fiber-forming proteins in modulating protein aggregation processes and their implications for therapeutic strategies against neurodegenerative diseases.

## Introduction

Fibrillar protein aggregation^1–3^ (so-called fibrillar self-assembly) plays a paradoxical role in biology, contributing both to disease^4^ and functional biomaterial formation.^5–7^ Amyloidogenic type of protein aggregation^8,9^ refers to the process by which certain proteins misfold and self-assemble^10–12^ into highly ordered, β-sheet-rich nanofibrillar structures known as amyloid fibrils.^13,14^ While amyloid fibrils formed by proteins like α-synuclein^15,16^ are closely linked to neurodegenerative disorders such as Parkinson’s disease,^17^ other fibrillar proteins, like silk fibroin,^18^ serve as crucial structural materials in nature.^19–21^ The interactions between these distinct classes of fiber-forming proteins^16,22,23^ remain an area of active investigation.

Amyloid fibrils are characterized by their β-sheet-rich structures,^24,25^ which make them highly stable and resistant to degradation.^26^ The process of amyloid formation typically involves a nucleation-dependent mechanism,^27^ where soluble monomeric proteins undergo conformational changes, leading to the formation of oligomers, which is promoted by intermolecular interactions,^28^ and further elongation into protofibrils, and ultimately mature fibrils.^29,30^ For example, under physiological conditions^31^, α-synuclein exists as an intrinsically disordered monomer, but under destabilizing conditions,^32^ it undergoes misfolding and self-assembly into β-sheet-rich amyloid fibrils.^33^ These fibrils accumulate in Lewy bodies,^34^ a hallmark of Parkinson’s disease, and are implicated in neurotoxicity.^28,35^

Silk fibroin (SF)^36^ is a mostly unstructured, high molecular weight (∼450 kDa) protein and its fibrillation is a highly controlled self-assembly process. Similarly to amyloids, SF fibrillation results in the formation of stable, β-sheet-rich structures, essential for the mechanical strength and flexibility of silk fibers.^37^ However, unlike amyloid aggregation, SF undergoes fibrillation through a regulated pathway that ensures the formation of biocompatible fibers without triggering cytotoxic effects.^38,39^ Furthermore, SF-based fibrils can be enzymatically degraded under specific conditions, facilitating their dynamic remodelling in biological systems. As a large, fibrous protein, SF can create a crowded molecular environment^40^ that potentially might affect the behaviour of other proteins in solution.

In the context of amyloid fibrillation, crowders, such as macromolecular proteins,^41^ osmolytes,^42^ or synthetic polymers,^43^ alter the kinetics and pathways of amyloid aggregation by modulating protein stability, solubility, and intermolecular interactions.^43–45^ Some crowders accelerate amyloid formation by increasing effective protein concentration via excluded volume effect,^41^ while others, depending on their nature and interactions, can stabilize intermediate states or even inhibit fibril formation.^46^ Understanding the exact effects of molecular crowding on protein aggregation is crucial for developing therapeutic strategies aimed at preventing pathological amyloid formation. In this study, we present a surprising counterexample: SF inhibits the aggregation of α-synuclein. Through structural and kinetic analyses, we explore how SF alters the aggregation dynamics of α-synuclein, influencing its self-assembly, rheological behavior of the protein-rich fluid, and susceptibility to enzymatic degradation. These findings highlight the complex interplay between fiber-forming proteins and suggest new avenues for modulating amyloid aggregation, with potential implications for neurodegenerative disease therapies.^47^

## Results and Discussion

### SF and α-synuclein protein: binding interaction and morphological changes

To test the effect of SF on the aggregation pathway of α-synuclein, we first analyzed the fibrillation of α-synuclein and SF separately, as well as in their mixture, under physiological conditions (37°C, 50 mM Tris, 150 mM NaCl, pH 7.4) (see *Methods*).

We first performed morphological analyses using transmission electron microscopy (TEM) and scanning electron microscopy (SEM) to visualize aggregation behaviour of α-synuclein and SF proteins. Our observations revealed that monomeric α-synuclein tends to form small (∼30 nm) agglomerates, likely resulting from its interaction with substrate surface. Aggregated α-synuclein formed oriented, needle-like nanofibrils (see **Figure 1a**).

**Figure 1.**
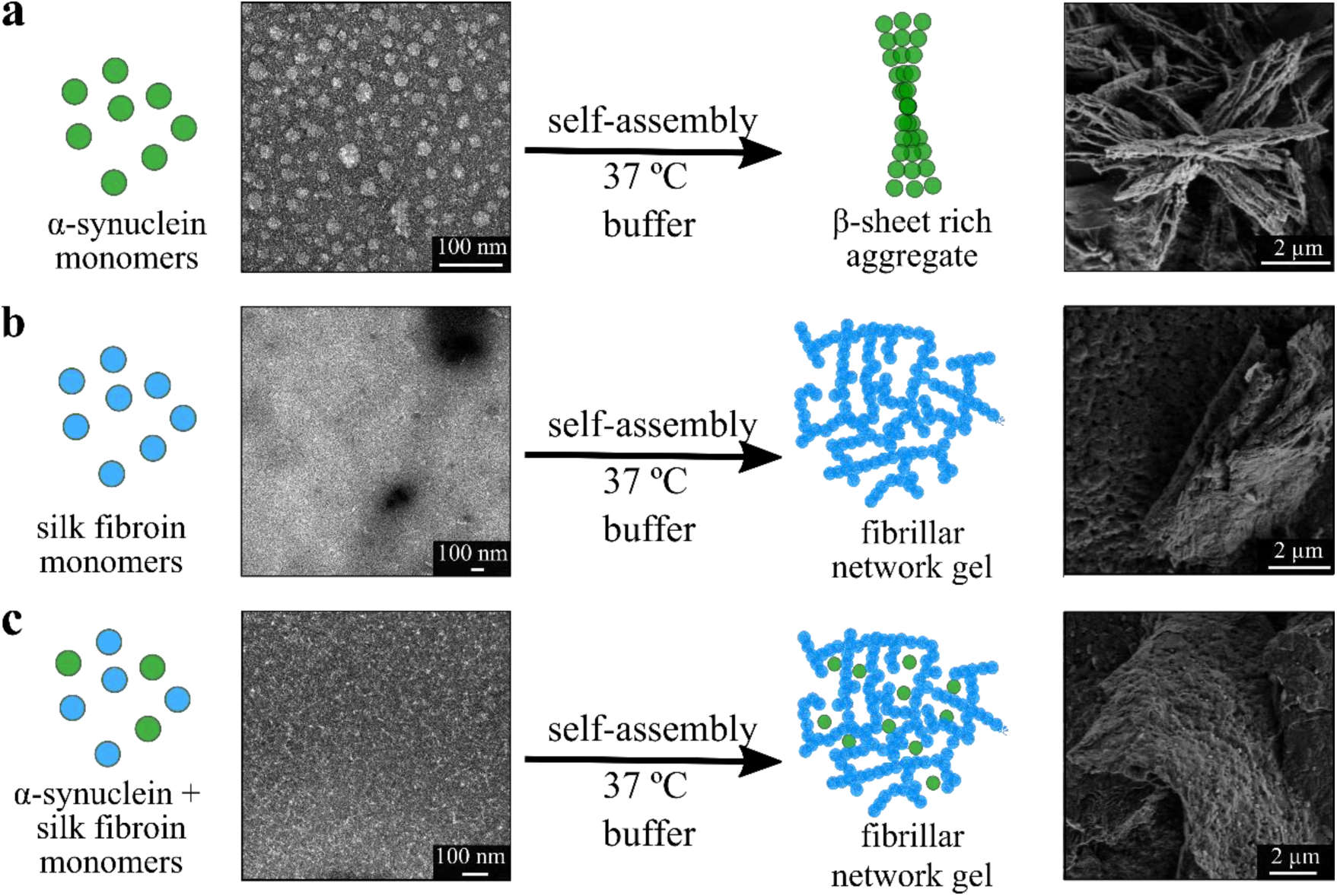
SF – α-synuclein co-assembly. **a**) Graphical representation supported by TEM (left) and SEM (right) images of α-synuclein fibrillation from monomeric to fibrillar state. **b**) Graphical representation supported by TEM (left) and SEM (right) images of silk fibroin (SF) fibrillation from monomeric to fibrillar state. **c**) Graphical representation supported by TEM (left) and SEM (right) images of SF- α-synuclein fibrillation from monomeric to fibrillar state. Scale bars for the SEM images are shown in bottom right corner.

Monomeric SF protein showed no signs of surface-induced aggregation. Aggregated SF developed into long, winding nanofibrils that interwined to form network-like, gel-like assemblies characteristic of SF self-organization (see **Figure 1b**). These observations are consistent with previous literature reports.^48^ Interestingly, the mixture of the two proteins closely resembled the morphological organization of SF only assemblies (see **Figure 1c)**, suggesting that SF may stabilize either the monomeric form of α-synuclein protein or facilitate the formation of nanofibrillar SF- α-synuclein co-aggregates.

To validate this hypothesis, we monitored changes in particle size distribution by using dynamic light scattering (DLS) technique.^49^ DLS analysis of non-aggregated monomeric proteins exhibited a good agreement between the theoretical^50^ and measured protein particle diameters^37^ (see **supplementary Figure S1a-c** and **TEM** of **Figure 1a**). DLS reports the hydrodynamic diameter— the apparent size of a molecule as it diffuses in solution, which depends not only on its molecular weight but also on its shape, hydration shell, flexibility, and solvation dynamics. Specifically, monomeric α-synuclein protein, with molecular weight (MW) of 14 KDa,^51^ displayed a measured DLS peak at ∼4.5 nm, with a negligible presence of larger aggregates ∼150 nm. Non-aggregated SF, with MW ∼450 KDa,^52^ the measured DLS peak appeared at ∼8 nm. Although α-synuclein and SF differ substantially in molecular weight, their measured DLS peaks do not scale proportionally with this difference. α-synuclein, despite its relatively low molecular weight, lacks a compact tertiary structure and behaves as an intrinsically disordered, expanded random coil in solution. Due to its flexible conformation and extensive hydration layer, DLS typically detects apparent hydrodynamic diameters of 3–5 nm for monomeric α-synuclein, consistent with reported literature values (Rh ≈ 2.8–4.5 nm).^53^ In contrast, SF is a high–molecular-weight, semi-structured protein composed of repetitive hydrophobic and β-sheet–forming domains separated by flexible amorphous segments. In aqueous environments, SF often adopts a collapsed, partially folded conformation, forming compact micellar or wormlike structures through intra- and intermolecular hydrophobic interactions.^54^ These dense, partially dehydrated assemblies result in a measured hydrodynamic diameter of approximately 7–10 nm—significantly smaller than would be expected for an unfolded linear polymer of comparable molecular weight. When two proteins were mixed in their monomeric state, the measured DLS size distribution showed an overlap between the two individual distributions measured for one component protein solutions, with two main peaks at ∼4.5 nm and ∼8 nm. The results indicate that when the two proteins are mixed in their monomeric forms, SF does not trigger α-synuclein aggregation. This is an unexpected observation since SF is a high-molecular-weight biopolymer and, like other similar biopolymers, is expected to impose spatial constraints on the amyloidogenic α-synuclein, creating a crowded environment.^40^ Under such crowded conditions, volume exclusion typically increases the effective concentration of the amyloidogenic protein, promoting and accelerating it’s aggregation.^55^ However, aggregation of these proteins in the presence of biopolymeric crowders can sometimes be also inhibited due to non-specific interactions^56^ between the crowder and amyloidogenic protein-in our case of study its SF (crowder) and α-synuclein (amyloidogenic protein).

To determine whether SF non-specifically interacts with α-synuclein and potentially stabilizes its monomeric form, we conducted micro-scale thermophoresis (MST)^57,58^ and enzyme-linked immunoassay (ELISA)^59^ analyses, and the results are summarized in **supplementary Figure S2**. Prior to MST and ELISA analyses the molecular weight of α-synuclein protein has been determined by using gel-electrophoresis (see **supplementary Figure S2a**). Briefly, MST measures the movement of a fluorescently labelled protein within a microscopic temperature gradient created by an infrared laser. This thermophoretic movement is affected by changes in protein size, charge, and interactions. To this end SF has been labelled with DyLight 650 (excitation at 652 nm; emission at 672 nm) via the NHS ester group (see *Methods*). When SF mixed with α-synuclein, binding can alter above-mentioned protein characteristics, thus changing the diffusion. Analysing these changes across different concentrations (**supplementary Figure S2b**) enabled us to determine the binding affinity, which was quantified as a Kd = 24 ± 17 nM —indicative of a relatively strong interaction^60^ (between SF and α-synuclein), comparable to high-affinity receptor-ligand bindings in biology. The ELISA assay assessed the protein interaction by measuring fluorescence intensity of labelled α-synuclein on a SF substrate. The increase in fluorescence signal (**supplementary Figure S2c**) with rising α-synuclein concentration supports the occurrence of interaction. Thus, results from both MST and ELISA analyses indicate that SF is capable of interacting with α-synuclein. However, the relatively high apparent affinity likely arises from non-specific or multivalent interactions rather than a defined binding interface. Given the intrinsically disordered and high–molecular-weight nature of SF, such interactions may involve transient electrostatic or hydrophobic contacts distributed along the protein chain. These weak, non-specific associations could nonetheless perturb α-synuclein’s conformational ensemble, influencing its stability, and aggregation behavior. Building on these findings, changes in the morphology of α-synuclein assemblies (formation of needle-like structures) over time of 4, 7, and 11 days we also visualized using confocal microscopy. The results, summarized in **supplementary Figures S3 and S4,** demonstrate the dimensional growth of α-synuclein structures in different media (water and buffer). These morphological changes provide further insights into the dynamic aggregation process of α-synuclein alone and in the presence of SF.

### Concentration and temperature factors– dominance of SF fibrillation over α-synuclein

We further investigated how different physico-chemical conditions affect SF–α-synuclein interactions and their aggregation behavior (see **Figure 2a-l**). Our analysis aimed to assess the extent to which SF can suppress α-synuclein fibril formation. To do this, we systematically decreased the concentration of SF, from SF:α-synuclein 1:1 to 1:10, 1:100, and 1:1000 (**Figure 2a–c**). After 14 days of incubation at 37°C under physiological conditions (see *Methods*), we analyzed the resulting protein assemblies. Fluorescence microscopy, using the amyloid-binding dye thioflavin T (ThT),^61^ was first employed to detect the presence of β-sheet-rich fibrillar structures. ThT fluorescence increases and undergoes a red shift upon binding to amyloid fibrils, making it a reliable marker for detecting amyloid-like aggregation (see *Methods*). We further used confocal microscopy to characterize the morphology and dimensions of the resulting aggregates (**Figure 2e)**. In the control sample (**Figure 2a**) containing α-synuclein only, the fibrillar structures were typically of ∼40 μm in length (see summary of length distribution in **Figure 2e**). When SF was introduced at the lowest concentration (SF:α-synuclein = 1:1000, **Figure 2b**), the aggregation behavior of α-synuclein was not significantly altered. However, increasing the SF concentration to a 1:100 ratio (**Figure 2c**) resulted in a noticeable reduction in fibril length, with average sizes reduced to approximately 20 μm. At the higher SF concentration tested (1:10 ratio, **Figure 2d**), the α-synuclein assemblies were markedly smaller, with lengths less than 10 μm. Importantly, while the size of the α-synuclein fibrils decreased with increasing SF concentration (**Figure 2e**), the overall abundance of fibrillar structures remained relatively constant across the different conditions. This suggests that SF modulates the morphology and growth dynamics of α-synuclein assemblies—primarily limiting their elongation—without significantly affecting the nucleation rate or total number of aggregates formed.

**Figure 2.**
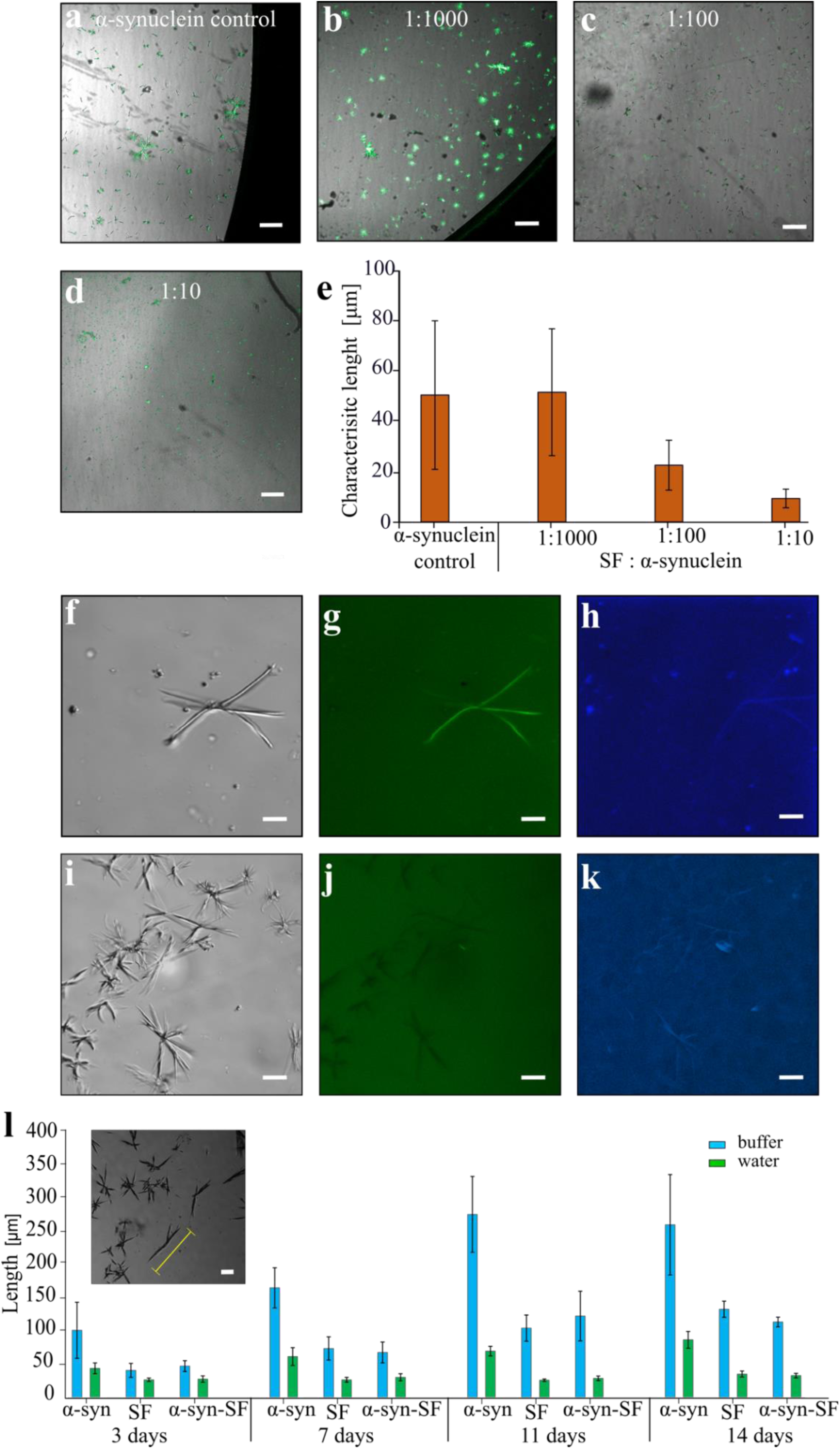
SF – α-synuclein assembly under varying environmental conditions: Merged light and fluorescent microscopy images of **a)** ThT-stained α-synuclein assembly; **b)** 1:1000 ThT-stained SF: α-synuclein assembly; **c)** 1:100 ThT-stained SF: α-synuclein assembly; **d)** 1:10 ThT-stained SF: α-synuclein assembly, after 11 days of, at 37°C, in buffer at a concentration of 10 μM. Scale bars for microscopy images are 20 μm. **e)** Bar chart summarizing the characteristic lengths of fibrillar structures calculated from confocal microscopy images. **f-h**) 1:1 ThT-stained SF: α-synuclein assembly and temperature (65°C) effect on self-assembly of α-synuclein, where **f)** showing brightfield (BF) image of α-synuclein assembly; **g)** confocal image of α-synuclein assembly stained with Alexa Fluor 568 dye; **h)** showing confocal (intrinsic fluorescence) image of α-synuclein assembly. **i-k**) The temperature (65°C) effect on self-assembly of SF:α-synuclein, where **i)** showing brightfield (BF) image of SF:α-synuclein assembly; **g)** confocal image of SF:α-synuclein assembly stained with Alexa Fluor 568 dye; **h)** showing confocal (intrinsic fluorescence) image of SF:α-synuclein assembly. Scale bars for microscopy images are 20 μm. **l)** Bar chart summarizing the changes in characteristic lengths of fibrillar structures calculated from confocal microscopy images. Insert showing light microscopy image of α-synuclein assemblies. Scale bar is 20 μm.

We next investigated how elevated temperatures influence the self-assembly behavior of α-synuclein, SF, and their mixtures. Exposing the proteins to the temperature of 65°C led to the formation of larger and more defined α-synuclein aggregates as shown in **supplementary Figures S5** and **S6**, consistent with thermally enhanced protein mobility and structural rearrangement.^62^ Interestingly, SF displayed distinct behavior under the same conditions: instead of forming a homogeneous gel-like matrix as observed at lower temperatures, it assembled into rigid, needle-like structures (see **supplementary Figures S5** and **S6**), suggesting a temperature-induced shift in its aggregation pathway.

To differentiate between α-synuclein and SF components within the mixed assemblies, we used fluorescently labeled α-synuclein (*see Methods*), tagged with Alexa Fluor 568 (excitation at 494 nm, emission at 517 nm). This labeling allowed us to visualize the distribution of α-synuclein within the composite structures using confocal microscopy (**Figure 2f-k**). When labeled α-synuclein was heated alone at 65°C (for 3 days), it retained a strong fluorescent signal, indicating successful assembly into fibrillar aggregates.

However, when co-incubated with SF under the same thermal conditions, the fluorescent signal was completely abolished, suggesting that α-synuclein was either excluded from the final assembly or its conformational state was altered to prevent fluorophore detection. This points to a dominant structural role of SF in the mixed assemblies formed at elevated temperature. Additionally, we observed an increase in the intrinsic fluorescence signal^63^ of the mixed structures compared to α-synuclein alone, particularly in the UV region (∼385 nm).The heightened fluorescence in the silk-containing structures thus supports the conclusion that silk dominates the assembly composition at 65°C. We also evaluated the evolution of these assembled structures over time in different solutions, namely buffer and water. Using ThT fluorescence and confocal microscopy, we monitored structural growth at 4, 7, 11, and 14 days (**Figure 2l** and **supplementary Figures S5** and **S6**). Size estimation, based on characteristic structural dimensions, revealed that both SF and SF–α-synuclein assemblies grew progressively until day 11, after which they plateaued (in water and buffer). Notably, there was no statistically significant difference in structural size between SF alone and the SF–α-synuclein mixtures, indicating that silk dominates the morphology and growth behavior of these structures even in the presence of α-synuclein.

### SF- α-synuclein aggregation mechanism

While the crowding effect induced by SF appears consistent across different concentration ratios and temperatures, we next focused on evaluating how SF influences the assembly of α-synuclein under physiological conditions (37°C in water and buffer). As illustrated in **Figure 3**, the assembly behavior followed two distinct patterns: one corresponding to α-synuclein alone, and the other encompassing SF and SF–α-synuclein mixtures. To monitor aggregation kinetics, we employed a ThT assay using 96-well plate (see *Methods*). Each well was loaded with 150 μL of sample containing 10 μM ThT. The kinetic traces, shown in **Figure 3a**, revealed an increase in fluorescence over long time scale (>10 days) for both SF and SF–α-synuclein mixtures, indicative of progressive fibril formation. Such kinetic profile with long time scale can be attributed to the relatively low SF concentration (10 μM), which requires extended incubation to achieve a fully developed fibrillar network. To describe the aggregation timeline more clearly, we defined two time points: t = 0, representing the starting condition, and t = SF-gel (indicative of completion of aggregative process, where t is >20 days), marking the formation of a SF-derived gel-like network, which denotes the completion of the aggregation process. This endpoint is primarily dictated by the intrinsic fibrillation kinetics of SF. Importantly, the α-synuclein-alone condition displayed minimal change in fluorescence intensity throughout the experiment, suggesting limited or slow fibril formation. In contrast, both the SF and SF–α-synuclein mixtures exhibited a substantial increase in fluorescence signal—approximately an order of magnitude higher than α-synuclein alone—confirming robust amyloid-like aggregation facilitated by SF.

**Figure 3.**
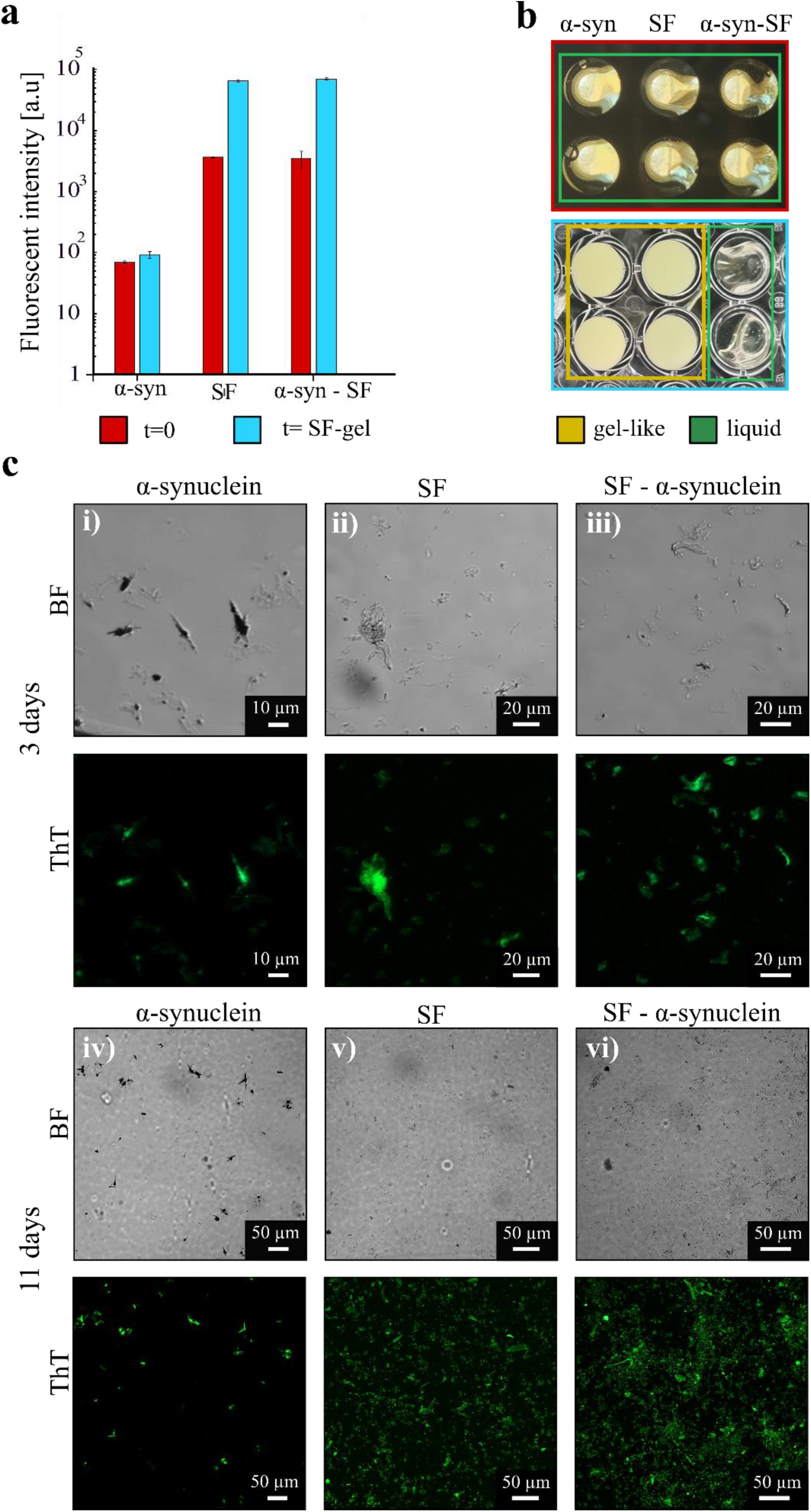
SF- α-synuclein aggregation mechanism. a) Chart showing fluorescent signal emitted from ThT stained assemblies of α-synuclein, SF and SF-α-synuclein fibrillation at t=0 and t=SF-gel, where t=SF-gel is the time when SF fibrillation is completed. b) pictures of 96-well plate depicting solutions of α-synuclein, SF and SF-α-synuclein at t=0 (top panel) and gelled solutions of α-synuclein, SF and SF-α-synuclein at t=SF-gel (bottom panel). c) Confocal images (bright field-top panel and green fluorescence bottom panel) of α-synuclein, SF and SF-α-synuclein assemblies acquired after 3 days (i) α-synuclein, ii) SF, iii) SF-α-synuclein) and 11 days ((iv) α-synuclein, v) SF, vi) SF-α-synuclein) under physiological conditions (see Methods). Scale bars are at the bottom right of each figure.

To visually corroborate the plate reader data, we imaged the wells before and after incubation as shown in **Figure 3b**. The SF and SF–α-synuclein samples transitioned into visibly gel-like structures by the end of the incubation period, while α-synuclein remained in a liquid state, forming isolated micron-sized aggregates, consistent with results shown in **Figures 1** and **3c** (panels ***i*** and ***iv***). These structures were detectable as early as day 3 by confocal microscopy and measured approximately <10 μm in length.

After 11 days of incubation, the α-synuclein assemblies had grown to around 30 μm and eventually reached their final size of ∼40 μm (as quantified in the bar chart in **Figure 2a**) by days 15–21. For SF and SF–α-synuclein samples, confocal microscopy revealed a similar progression: early-stage aggregates were visible after 3 days, and by day 11, the samples exhibited widespread and uniform gelation, indicative of extensive network formation (**Figure 3c, panels ii, v,** and **iii, vi**) prior to achieving the t=SF-gel point at which SF fibrillation is completed. These observations further support the role of SF in facilitating and possibly templating the aggregation process, either independently or in the presence of α-synuclein. In summary, under physiological conditions, SF promotes a distinct and robust aggregation pathway that overrides the slower and less extensive self-assembly of α-synuclein alone.

### Degradability of SF- α-synuclein fibrillar assemblies

We further analyzed and compared the degradability of the assembled structures of α-synuclein, SF, and the SF–α-synuclein mixture. To assess their susceptibility to enzymatic breakdown, we treated the samples with two proteolytic enzymes—trypsin (designated with “T” in legends of **Figure 4a**) and protease XIV (designated with “P” in legends of **Figure 4a**) —each at a concentration of 1 μM. At the initial stage, aggregated α-synuclein sample contained needle-like assemblies, whereas SF appeared as fibrillar network gel. The mixture of both proteins displayed a morphology closely resembling that of the SF fibrillar network gel. The degradation kinetics were monitored based on changes in ThT fluorescence measured by using plate reader (**Figure 4a)**. In the initial phase of enzymatic activity, SF showed the fastest and prolonged degradation, particularly under the activity of protease XIV, which continued to degrade the protein assemblies at a slower but steady rate for up to 72 hours. Trypsin, in contrast, exhibited a more rapid decline in activity (for SF assemblies) after the initial burst. Remarkably, both SF and SF–α-synuclein samples displayed similar degradation profiles, with progressive loss of fluorescence indicating breakdown of the aggregated structures. In stark contrast, α-synuclein alone remained largely unaffected by either of two enzymes, suggesting its well-known resistance to proteolysis due to the stable and compact β-sheet-rich fibrillar architecture of mature amyloid structures. These results were validated by confocal microscopy images acquired after 3 and 11 days of enzymatic treatment (**Figure 4b**). The α-synuclein assemblies retained their structural integrity and fluorescence over time (**Figure 4b** panels ***i*** and ***iv***). Conversely, SF and SF–α-synuclein samples showed pronounced degradation (**Figure 4b** panels ***ii, v*** and ***iii, vi***). After one day of exposure, the aggregated networks were visibly compromised, with patchy fluorescence signals and disrupted structures. By day 6, the fluorescence signal had completely disappeared, indicating extensive or complete enzymatic digestion.

**Figure 4.**
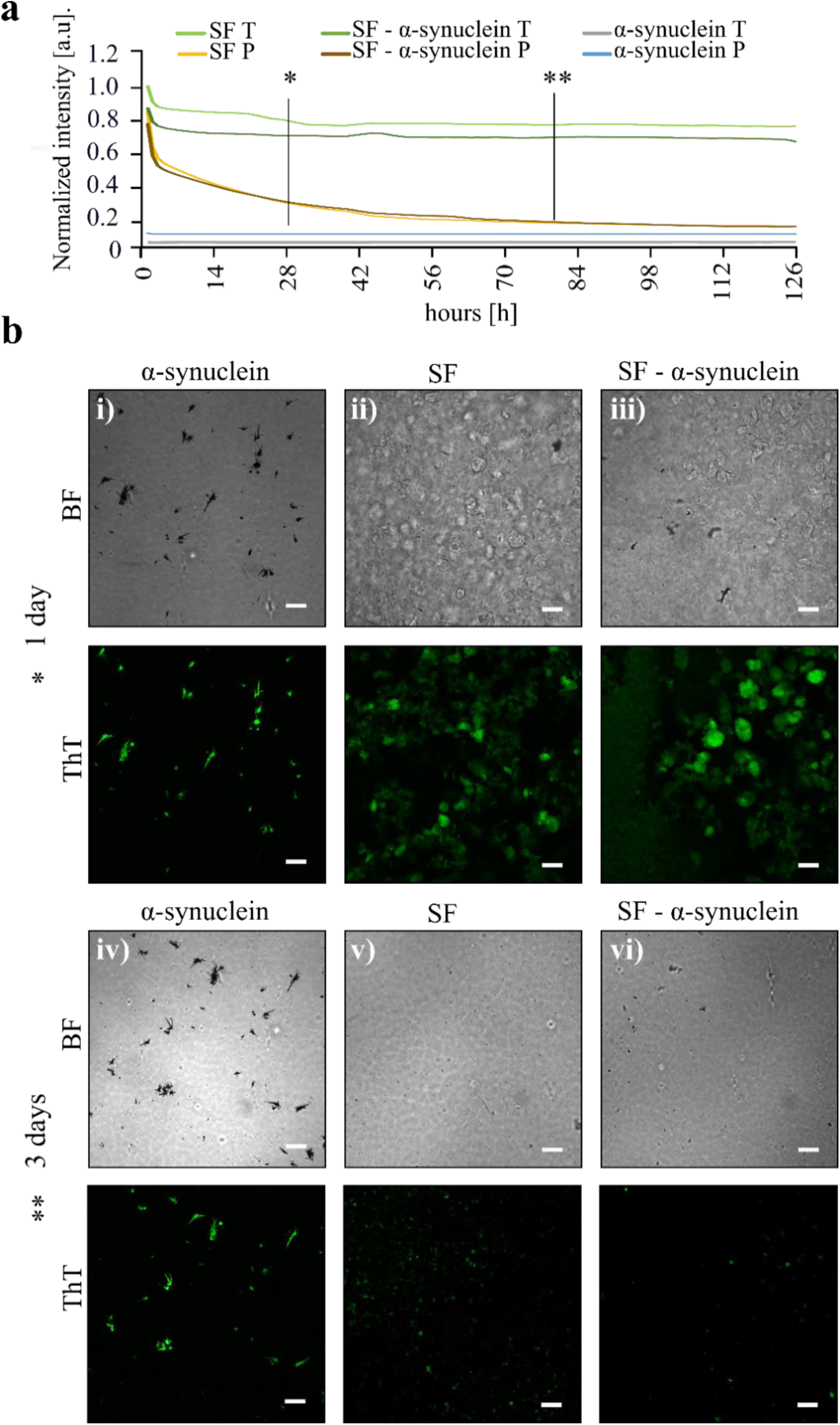
Degradation of α-synuclein SF and SF-α-syn assemblies. **a**) Graph summarizing changes in ThT fluorescent signal over time emitted from stained α-synuclein, SF, and SF-α-synuclein assemblies exposed to trypsin and protease XIV enzymes. **b**) Confocal (bright field-top panel, fluorescence-bottom panel) images of (i) α-synuclein, (ii) SF, (iii) SF – α-synuclein assemblies enzymatically treated for 1 day and (iv) α-synuclein, (v) SF, (iv) SF – α-synuclein-treated for 3 days long. **c**) Confocal images acquired after 3 days. Scale bars are 50 μm.

### Structural changes in SF- α-synuclein assemblies

Protein assembly is a dynamic, multistep process that often follows distinct kinetic pathways, involving the structural reorganization of molecules that ultimately influences the physical and mechanical properties of the resulting material. To gain insight into this relationship, we investigated the mechanical behavior of α-synuclein, SF, and SF–α-synuclein mixtures over time using oscillatory rheology measurements.^64,65^ Oscillatory rheology analysis was performed using a time sweep protocol (*see Methods*) to monitor changes in the storage modulus (G′) at 37°C. The storage modulus represents the elastic component of a material’s viscoelastic behavior,^66^ and its increase over time under constant strain and frequency is indicative of gelation or structural maturation. As shown in the rheological data (**Figure 5a**), α-synuclein alone exhibited a relatively steady G′ over time, suggesting minimal structural evolution and the absence of a transition to a gel-like state. In contrast, both SF and SF–α-synuclein samples showed a marked increase in G′ between 10 and 14 hours of incubation, indicating a clear transition toward a more rigid, networked structure. This abrupt rise in mechanical stiffness reflects the onset of gelation or fibrillar network formation, consistent with the aggregation patterns observed earlier (see **Figure 3a–b)**. Triplicate measurements (**supplementary Figure S7**) confirmed the reproducibility of these trends. Interestingly, the observed aggregation and gelation occurred more rapidly than in the ThT-assay. For example, while for ThT-assay the aggregation kinetics lasts for > 20 days, in the oscillatory rheology analysis the result is achieved > 16 hours. This difference can be attributed to the distinct geometrical configurations of the experimental setups. The rheological analysis employed a sample volume of 170 μL with a radius of 0.02 m, yielding a surface-to-volume ratio of approximately 0.0375. In comparison, the 96-well plate used for fluorescence assays had a smaller radius (0.0025 m) and volume (150 μL), resulting in a much lower surface-to-volume ratio of 0.34. Thus, the setup for rheological analysis offers a surface-to-volume ratio roughly 54 times higher than that of the plate reader-based kinetic analysis, promoting more efficient heat transfer and thus, accelerates aggregation kinetics. To ensure environmental consistency, all rheological measurements were conducted in a sealed chamber with controlled temperature and humidity.

**Figure 5.**
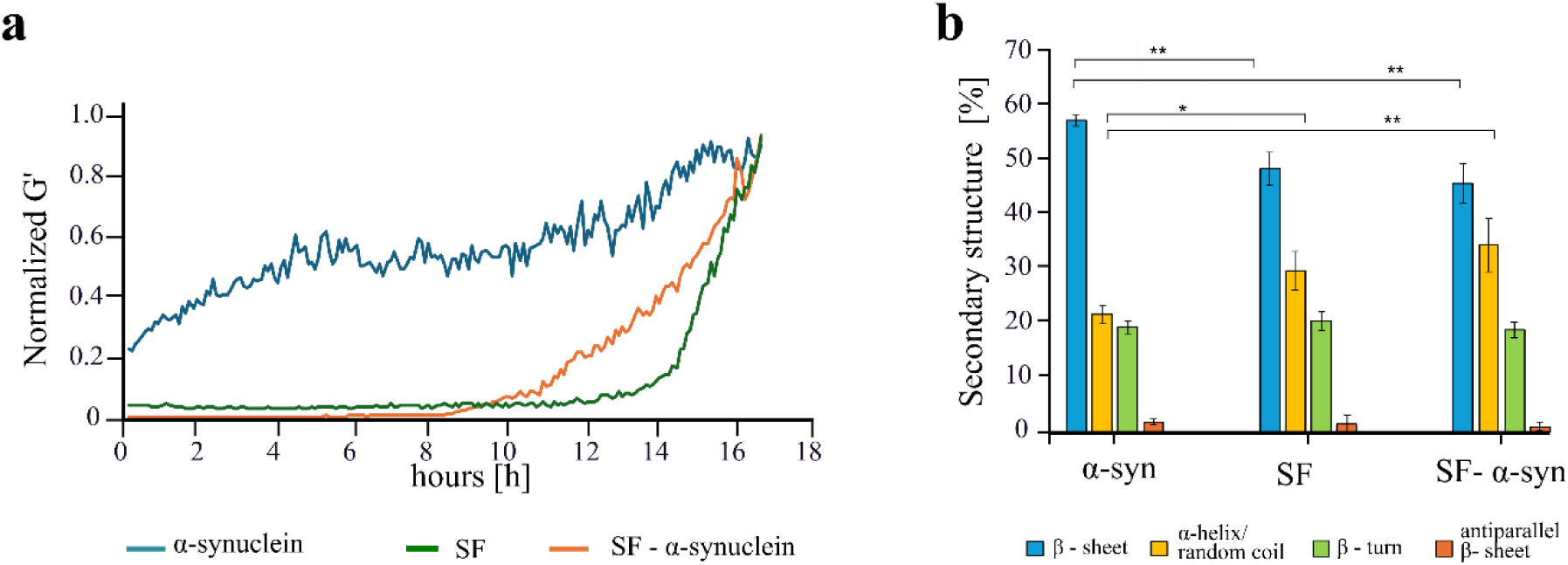
Kinetic and structural analysis of α-synuclein, SF, and SF – α-synuclein assemblies. Graph depicting changes in storage modulus (G’) over time for α-synuclein, SF, and SF – α-synuclein solutions, tested by rheological analysis. **b**) Chart summarizing changes in secondary structure of α-synuclein, SF, and SF – α-synuclein assemblies.

To further understand the molecular rearrangements underlying the observed mechanical transitions, we performed secondary structural analysis using Fourier Transform Infrared Spectroscopy (FTIR) following thermal treatment. **Figure 5b** shows the FTIR spectra deconvoluted in the Amide I region (1600–1700 cm⁻¹), which is highly sensitive to protein secondary structures. This region was analyzed to quantify different structural motifs: β-sheets (1600–1635 cm⁻¹), α-helix and random coil (1635–1665 cm⁻¹), β-turns (1665–1690 cm⁻¹), and antiparallel β-sheets (1690–1705 cm⁻¹), as detailed in **supplementary Figure S8.** The deconvoluted spectra revealed distinct structural differences between assembled/fibrillar state of α-synuclein and SF and SF-α-synuclein samples. α-synuclein assemblies displayed a lower proportion of α-helix and random coil structures, whereas SF and SF–α-synuclein exhibited low content of β-sheets—hallmarks of silk’s structural organization and amyloid-like fibrillation. Statistical analysis using ANOVA t-tests confirmed these differences: the β-sheet content differed significantly (p < 0.01) between α-synuclein and both SF and SF–α-synuclein. Similarly, the α-helix/random coil content differed with statistical significance (p < 0.05 for α-synuclein vs. SF and p < 0.01 for α-synuclein vs. SF–α-synuclein). These findings further validate that SF induces conformational changes in α-synuclein assemblies, favoring lower content of ordered β-sheet structures. **Supplementary Figure S8** also presents normalized overlapping FTIR spectra, demonstrating how variations in raw data influence the deconvolution process. For instance, in the α-helix/random coil region, α-synuclein displays a broad, indistinct peak, while its signal in the native β-sheet range is relatively narrow—reflecting a more ordered structure compared to SF-based assemblies. Notably, α-synuclein secondary structures were also examined using circular dichroism (CD) in solution and compared with FTIR data of lyophilized samples (see **supplementary Figure S9**). Indeed, while in monomeric form, α-synuclein showed a dominant α-helix/random coil secondary structure, whereas the initial lyophilized conditions presented a sharp increase of β-sheet content. This dual approach highlights how the structural conformation of α-synuclein varies depending on environmental conditions^67,68^, supporting previous findings on the context-dependent polymorphism of amyloid fibrils

## Conclusion

Fibrillar, β-sheet-rich, type of protein self-assembly can trigger either formation of functional material or disease-associated constructs. In this study, we demonstrate that SF can inhibit the aggregation of the neurodegeneration-associated protein α-synuclein. Using MST and ELISA analyses along with structural characterization and oscillatory rheology studies, we confirmed that SF alters the structural transitions and self-assembly of α-synuclein through macromolecular crowding and non-specific interactions, ultimately supressing its amyloidogenic aggregation. We then analyzed SF and α-synuclein co-assembly path under varying conditions, including buffer, temperature and concentration, by using kinetics assays and microscopy characterization. Thus, under physiological conditions, SF significantly altered α-synuclein aggregation by suppressing β-sheet-rich fibrils formation and promoting a uniform nanofibrillar gel network. Enzymatic degradation tests showed complete dissolution of SF and SF–α-synuclein composites, while stable α-synuclein fibrils remained resistant. This enhanced degradability of the SF-containing assemblies highlights their potential physiological relevance, suggesting that incorporating functional, biocompatible proteins such as silk may facilitate natural clearance or remodeling of otherwise persistent amyloid aggregates in biological environments. These findings reveal that SF interfere with α-synuclein both molecularly and structurally, effectively inhibiting amyloidogenic filament formation. This positions silk protein as a promising candidate for therapeutic strategies targeting neurodegenerative diseases.

## Methods

### α-synuclein purification

BL21 (DE3) cells transformed with the pT7-7 plasmid encoding wild-type (WT) α-synuclein were grown in LB medium at 37°C. Protein expression was induced with 1 mM IPTG (isopropyl β-D-1-thiogalactopyranoside) when the culture reached an OD₆₀₀ of 0.4– 0.6, and incubation continued for an additional 4 hours at 37°C. Induced bacterial cultures were pelleted and lysed by sonication. The lysate was centrifuged at 18,000 × g for 20 minutes, and the resulting supernatant was boiled for 20 minutes, then centrifuged again under the same conditions. The final supernatant was subjected to purification by anion exchange chromatography (HiPrep 16/10 Q FF, Cytiva), followed by gel-filtration chromatography using a HiLoad 16/600 Superdex 200 column (Cytiva). α-synuclein was then collected, lyophilized and stored at −20 °C. Before experiment, concentration and dilution (buffer (50 mM Tris, 150 mM NaCl, pH 7.4) or water (milliQ)) of lyophilized α-synuclein, was performed.

### Silk Fibroin

*Bombyx mori* (*B.mori*) silkworm cocoons were peeled to obtain a total weight of 5 grams of silk fibroin (SF) protein. Two identical solutions were prepared by dissolving 20 mM (4.24 g) of sodium carbonate (>99.5%, Fisher Chemical, USA) in 2 L of Milli-Q water and heated to 95 °C to chemically remove the sericin coating surrounding the fibroin fibers. The cocoon material was immersed in the boiling solution for 15 minutes, twice, then thoroughly rinsed with Milli-Q water and air-dried. The dried, sericin-free fibers were then dissolved in a 9.3 M lithium bromide (LiBr) solution, using a volume ratio of 1:4 (1 g of silk in 4 mL of LiBr), at 65 °C for 4 hours. The resulting solution was transferred into a dialysis membrane (10 kDa MWCO) and dialyzed against 2 L of Milli-Q water for 3 days, with the water being changed three times per day. After dialysis, the solution was centrifuged at 12,700 × *g* at 4 °C for 20 minutes, twice, to remove insoluble residues. The final SF solution was stored at 4 °C for subsequent experimental use. Adaptation of concentrations and dilution tests were further performed using α-synuclein protein.

### Protein – Protein Interaction

To measure the binding constant, two quantitative techniques were employed. The first technique was MicroScale Thermophoresis (MST) (NanoTemper Technologies, Germany). Briefly, the protein sequences were obtained from the UniProt database.^69^ The extinction coefficients were then calculated using the ExPASy ProtParam tool.^70^ The values used were 5,960 M⁻¹·cm⁻¹ for α-synuclein and 26,025 M⁻¹·cm⁻¹ for reconstituted silk fibroin (RSF) at 280 nm. Protein concentrations were subsequently determined using spectrophotometer (*NanoDrop One*, Thermo Fisher Scientific) by applying Beer– Lambert’s law, after background subtraction for wavelengths greater than 350 nm:

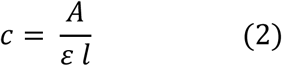

being A, ε, l, c, absorption [ODI], molar extinction coefficient 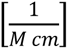 length [*cm*] and concentration [*M*]of solution, respectively. After diluting the stock solutions to 10 μM α-synuclein and 10 nM RSF, the RSF was labeled with DyLight 650-NHS Ester (Thermo Fisher Scientific) at a final concentration of 30 μM. Labelling was performed by incubating the RSF with the fluorophore for 1 hour at room temperature in 0.2 M sodium carbonate buffer (pH 8.4), protected from light.

For the **MST assay**, α-synuclein and labelled RSF were mixed in 16 different molar ratios, with α-synuclein undergoing a serial two-fold dilution across the samples. Specifically, 10 μL of buffer (or water) was added to each of 16 tubes (200 μL capacity). Then, 10 μL of the 10 μM α-synuclein solution was added to the first tube and mixed thoroughly. From this, 10 μL was transferred to the second tube, mixed, and so on—serially diluting the α-synuclein concentration two-fold in each step. The remaining 10 μL in the final tube was discarded. After the dilution series was completed, 10 μL of 10 nM labeled RSF was added to each tube to maintain a constant RSF concentration across all samples. The 16 tubes were then centrifuged at 14,400 rpm for 10 minutes at 4°C. Each sample was loaded into capillaries (MO-K022, NanoTemper), following the order of decreasing α-synuclein concentration, and analyzed using MST to determine the binding affinity.

The second technique used to assess the interaction between α-synuclein and RSF was **enzyme-linked immunosorbent assay (ELISA)**. For ELISA, lyophilized α-synuclein was -dissolved in Milli-Q water to a concentration of 2 mg/mL and labeled with Alexa Fluor 568 NHS Ester (Sigma, Cat. No. 20003). The fluorophore was first dissolved in 15 μL of DMSO at a concentration three times higher than that of the α-synuclein solution, then mixed with α-synuclein in a 2 kDa MWCO dialysis bag. The mixture was incubated for 1 hour at room temperature in 0.2 M NaHCO₃ buffer (pH 8.3). After labeling, the dialysis bag was transferred to working buffer and dialyzed overnight at 4°C. For ELISA, Maxisorp 96-well plates (Sigma, Cat. No. CLS3896-48EA) were used. RSF protein, pre-dialyzed in 0.1 M Na₂CO₃ (pH 9) overnight, was added to each well at 100 μL/well with a concentration of 2 mg/mL, and the plate was incubated overnight at 4°C. The supernatant was then discarded. Next, 100 μL/well of blocking buffer (2% BSA in working buffer) was added and incubated for 1 hour at room temperature, followed by removal of the supernatant. A serial dilution of labeled α-synuclein ranging from 1 μg/mL to 0.1 ng/mL was added to the wells and incubated for 1 hour at room temperature. After incubation, the wells were rinsed twice with working buffer, and fluorescence was measured using a plate reader.

For microscopy detection, Alexa Fluor 488 NHS Ester (Sigma, Cat. No. A20000) was used to label α-synuclein following the same protocol described above.

### Optical microscopy

Fluorescence microscopy was employed to detect micrometer-scale structures formed in a 96-well plate using a confocal microscope (Carl Zeiss LSM 800 with Airyscan, Germany). Imaging was performed using Thioflavin-T (ThT)fluorescence with 488 nm excitation, intrinsic fluorescence (IF) with 405 nm excitation, and brightfield (BF) imaging. Plan-Apochromat 10× and 20× objective lenses (air) were used, along with an additional 1.6× software zoom (Optovar) for enhanced resolution.

### Transmission Electron Microscopy (TEM)

Fibrils and monomers were deposited onto freshly glow-discharged, 300-mesh Formvar carbon-coated TEM grids (Electron Microscopy Sciences, Biolyst) and negatively stained with 2% aqueous uranyl acetate (Electron Microscopy Sciences, Biolyst). Imaging was carried out using a FEI Tecnai T12 transmission electron microscope operated at 120 kV.

### Secondary Structure Analysis

**Fourier Transform Infrared Spectroscopy (FTIR)** was used to analyze the protein secondary structure within the amide I region (1595–1720 cm⁻¹). Measurements were performed using a Nicolet iS50 FT-IR spectrometer equipped with a mounted ATR Smart iTX accessory. At least three independent samples were analyzed. Spectra were collected in the range of 400–4000 cm⁻¹ with a resolution of 4 cm⁻¹, averaging 32 scans per measurement. Solvent and air backgrounds were collected separately and recursively subtracted from each sample spectrum. Post-processing was performed using OriginPro software. Data were extracted specifically from the amide I region (1595– 1720 cm⁻¹). Baseline subtraction was followed by normalization of the spectra. Seven Gaussian peaks were fitted at the following vibrational wavenumbers: 1609, 1621, 1631, 1650, 1673, 1695, and 1703 cm⁻¹. The fit was considered satisfactory when the chi-square tolerance was below 1 × 10⁻⁵. For quantitative assessment of secondary structure: 1609, 1621, and 1631 cm⁻¹ were assigned to intermolecular β-sheets, 1650 cm⁻¹ to α-helix and random coil, 1673 and 1703 cm⁻¹ to β-turns, and 1695 cm⁻¹ to antiparallel amyloid β-sheets.

**Circular Dichroism (CD)** spectra were acquired for diluted protein samples (1 μM) in buffer (37°C, 50 mM Tris, 150 mM NaCl, pH 7.4) the 85–250 nm range using a J-715 spectropolarimeter (Jasco, Tokyo, Japan) with a data resolution of 0.5 nm. Samples were measured in a 0.1 mm quartz cuvette. Post-processing and quantification of secondary structure content were carried out using the DichroWeb online platform, applying the Contin-LL algorithm for structural deconvolution.

### Degradation

α-Synuclein fibrillar assemblies were subjected to enzymatic degradation using trypsin (0.25%)–EDTA (0.02%) solution (sterile, Bio-Lab, Israel) and Protease XIV (Sigma, Cat. No. 9036-06-0). Enzymes were added to samples that had undergone thermal treatment for 14 days at 37 °C, at a final concentration of 1 μg/mL. The 96-well plate was then resealed and incubated for the degradation period (3 days in our case) at 37 °C.

### Rheological analysis

**A rheometer** (Discovery HR-2, TA Instruments) equipped with a cone geometry (0.5° angle) was used to monitor the changes in rheological behavior of the solution samples. The cone geometry was selected to ensure uniform shear distribution across the sample. Prior to each measurement, 180 μL of sample solution was carefully loaded onto the rheometer stage, and the cone-to-plate gap was set to 17 μm.

**Oscillatory rheology** was conducted using a time sweep mode, applying an oscillatory shear rate of 0.3 rad/s at a frequency of 1 Hz, maintained at a constant temperature of 37 °C. Measurements were recorded every 10 minutes over a total duration of 18 hours. To prevent sample evaporation and maintain stable environmental conditions, an insulating chamber was employed throughout the experiment.

## Acknowledgments

M.E.M. thanks the Sergio Lombroso Fellowship (for Cancer Research) for financial support. U.S. acknowledge financial support from NATO (G6194). This work was supported by a research grant from Human Frontiers Scientific Program-HFSP (https://doi.org/10.52044/HFSP.RGP0202024.pc.gr.194172). In addition, U.S. thanks the Perlman family for funding the Shimanovich Lab at the Weizmann Institute of Science: “This research was made possible in part by the generosity of the Harold Perlman Family.” The authors would like to acknowledge partial support from the GMJ Schmidt Minerva Center of Supramolecular Architectures at the Weizmann Institute. This research was supported by a research grant from the Tom and Mary Beck Center for Advanced and Intelligent Materials at the Weizmann Institute of Science, Rehovot, Israel. Y. B. is the incumbent holder of the Beatrice Barton Research Fellowship.

## Supplementary Figures

**Supplementary Figure S1.**
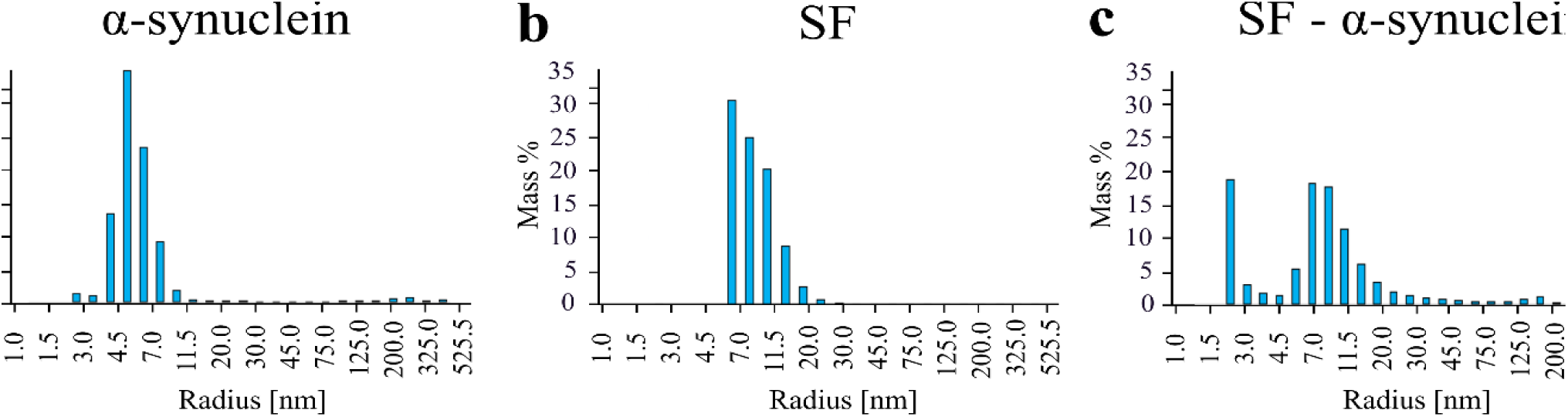
Dynamic Light Scattering (DLS) The acquisition was performed after diluting the proteins in buffer. **a**) DLS auto correlation function assessing the size distribution (by mass in %) for monomeric α-synuclein, with a peak value of ∼4.5 nm. **b**) DLS size distribution (by mass in %) for monomeric SF, with a peak value of ∼7.0 nm. **c**) DLS bimodal size distribution (by mass in %) for monomeric SF-α-synuclein, with peaks of ∼4.5 nm and of ∼8.0 nm.

**Supplementary Figure S2.**
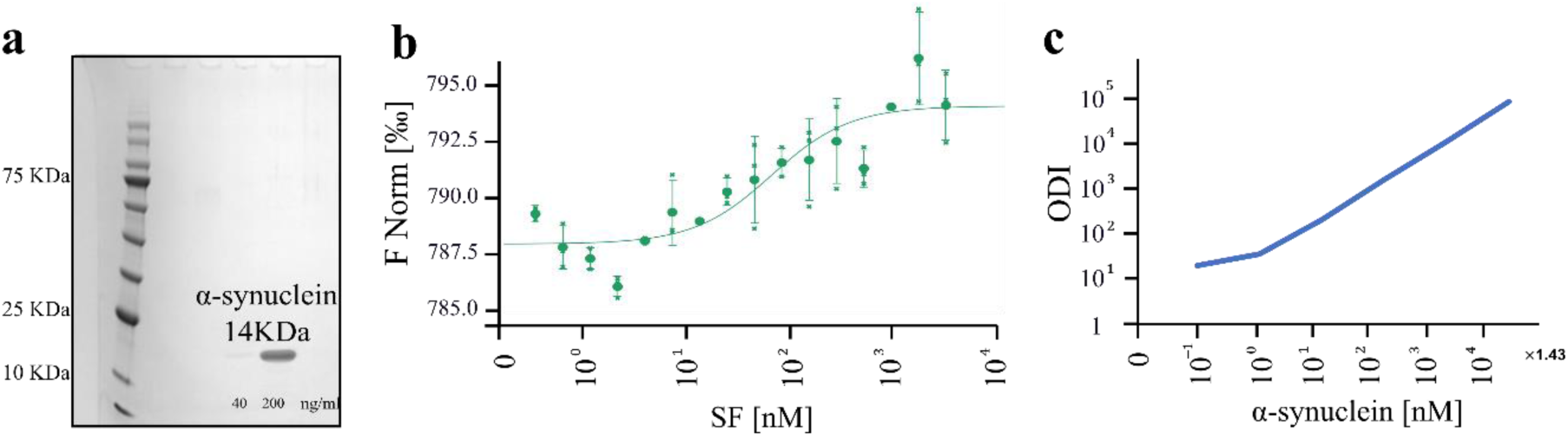
Characterization of protein-protein interactions. **a**) picture of the gel from gel-electrophoresis analysis, depicting band (14 kDa) of α-synuclein protein. **b**) Graph summarizes MicroScale Thermophoresis (MST) analysis confirming the interaction between the two proteins (stained α-synuclein with NHS dye and SF) with the affinity coefficient of Kd = 24 ± 17 nM. **c**) Graph summarizes enzyme-linked immunosorbent assay (ELISA) analysis, showing the interaction between SF and α-synuclein by staining SF with the fluorescent dye. Experiments conducted at room temperature.

**Supplementary Figure S3.**
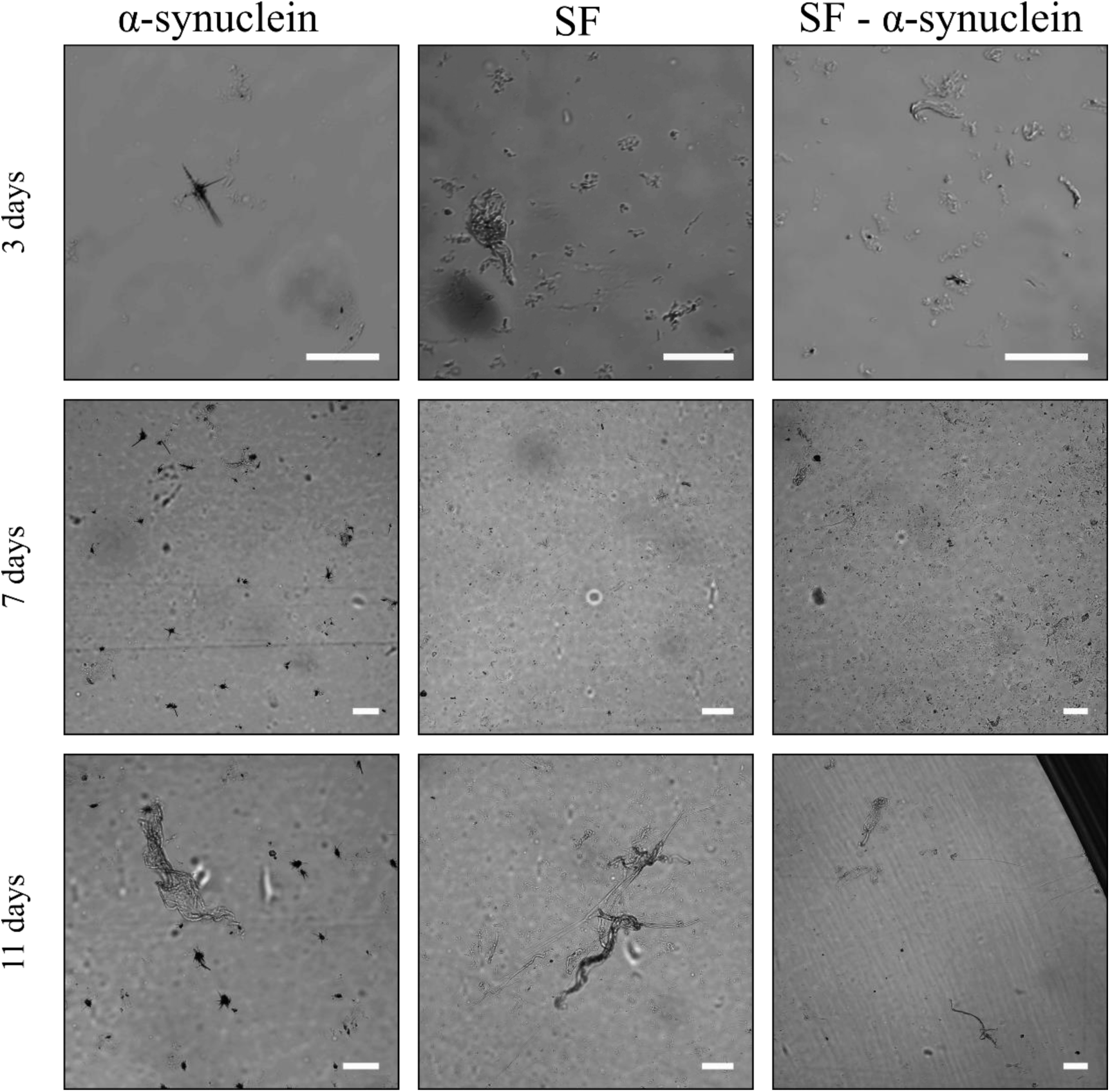
Aggregation over time of α-synuclein, SF and SF- α-synuclein. Light microscopy images of α-synuclein, SF and SF- α-synuclein assemblies formed within 3, 7 and 11 days in buffer at 37°C. Scale bars are 50 μm.

**Supplementary Figure S4.**
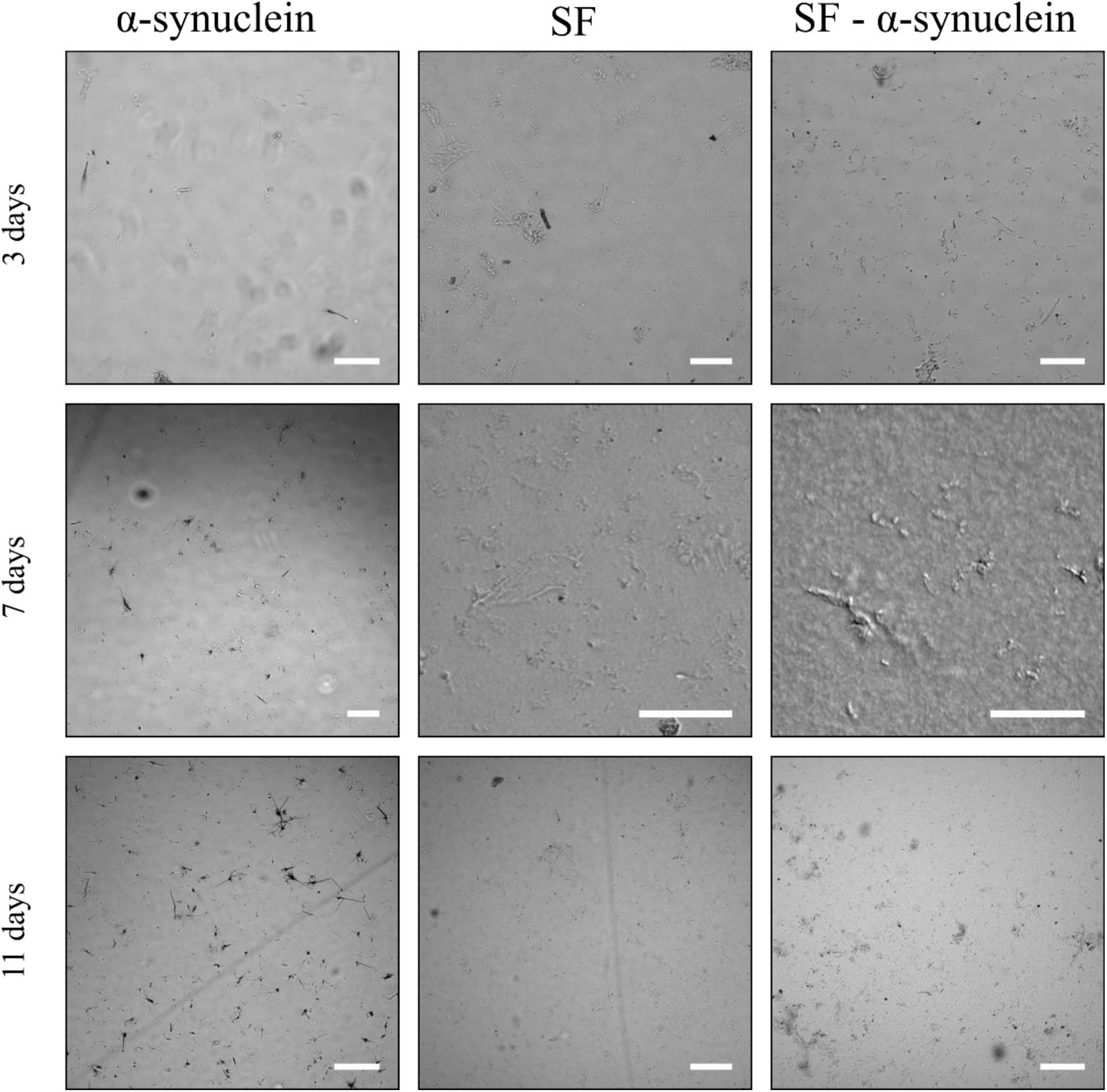
Aggregation over time of α-synuclein, SF and SF- α-synuclein. Light microscopy images of α-synuclein, SF and SF- α-synuclein assemblies formed within 3, 7 and 11 days in water at 37°C. Scale bars are 50 μm.

**Supplementary Figure S5.**
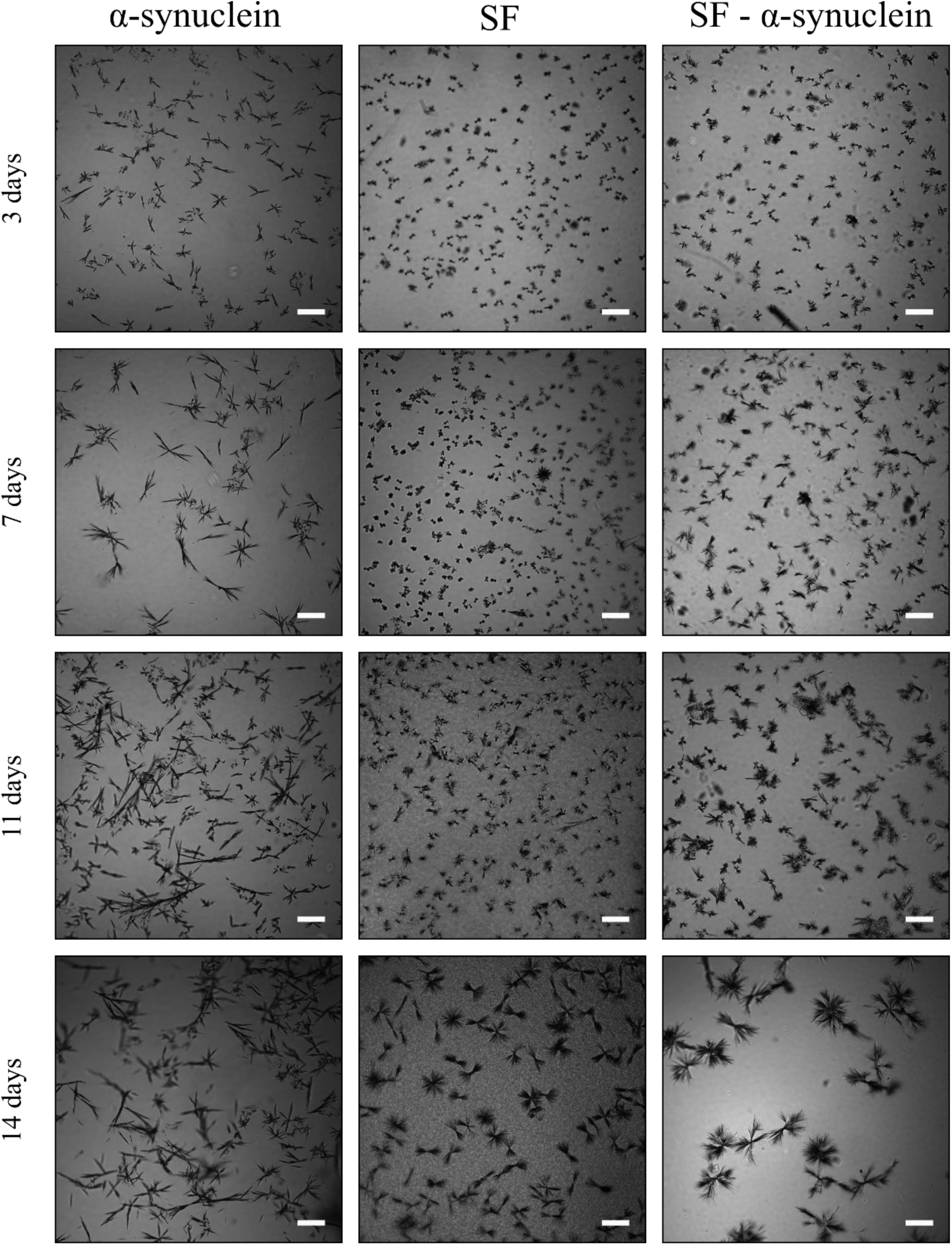
Aggregation over time of α-synuclein, SF and SF- α-synuclein. Light microscopy images of α-synuclein, SF and SF- α-synuclein assemblies formed within 3, 7 and 11 days in buffer at 65°C. Scale bars are 50 μm.

**Supplementary Figure S6.**
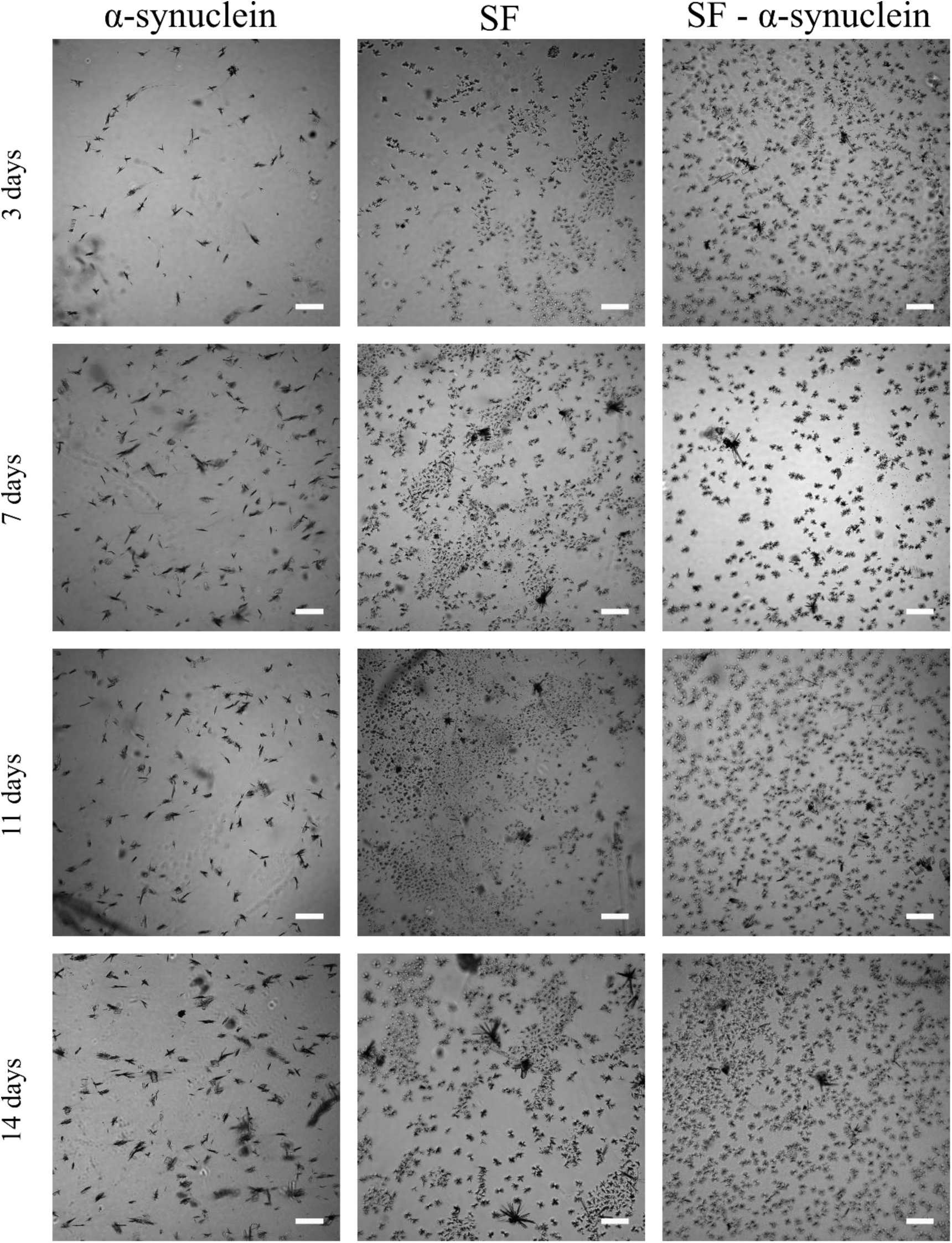
Aggregation over time of α-synuclein, SF and SF- α-synuclein. Light microscopy images of α-synuclein, SF and SF- α-synuclein assemblies formed within 3, 7 and 11 days in water at 65°C. Scale bars are 50 μm.

**Supplementary Figure S7.**
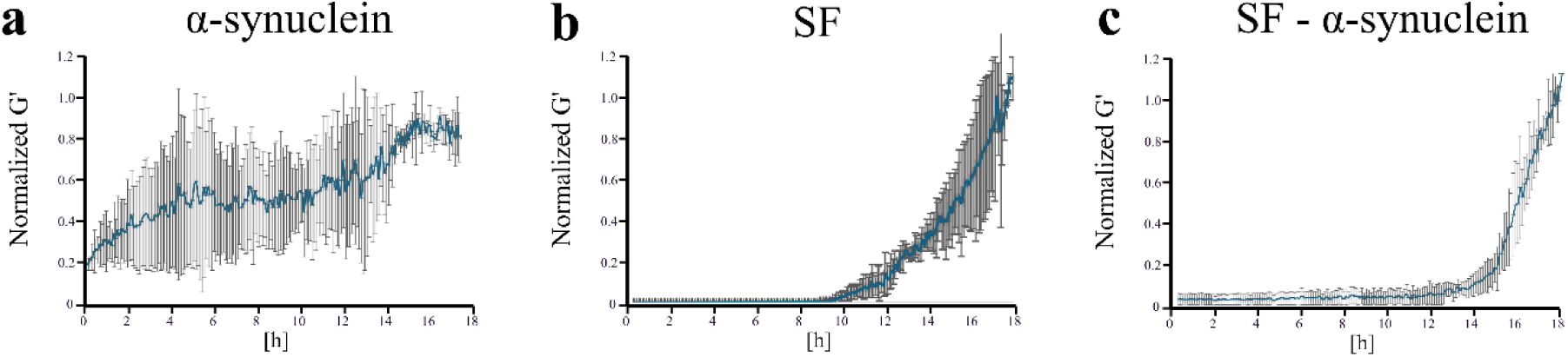
Oscillatory rheology. Graph summarizing results of oscillatory rheology analysis for **a)** α-synuclein, **b)** SF and **c)** SF- α-synuclein assemblies, revealing the changes in the storage modulus G’ at 37 °C. Experimental conditions included 1Hz frequency and 0.3 rad/s with 10 minutes interval for 18 hours.

**Supplementary Figure S8.**
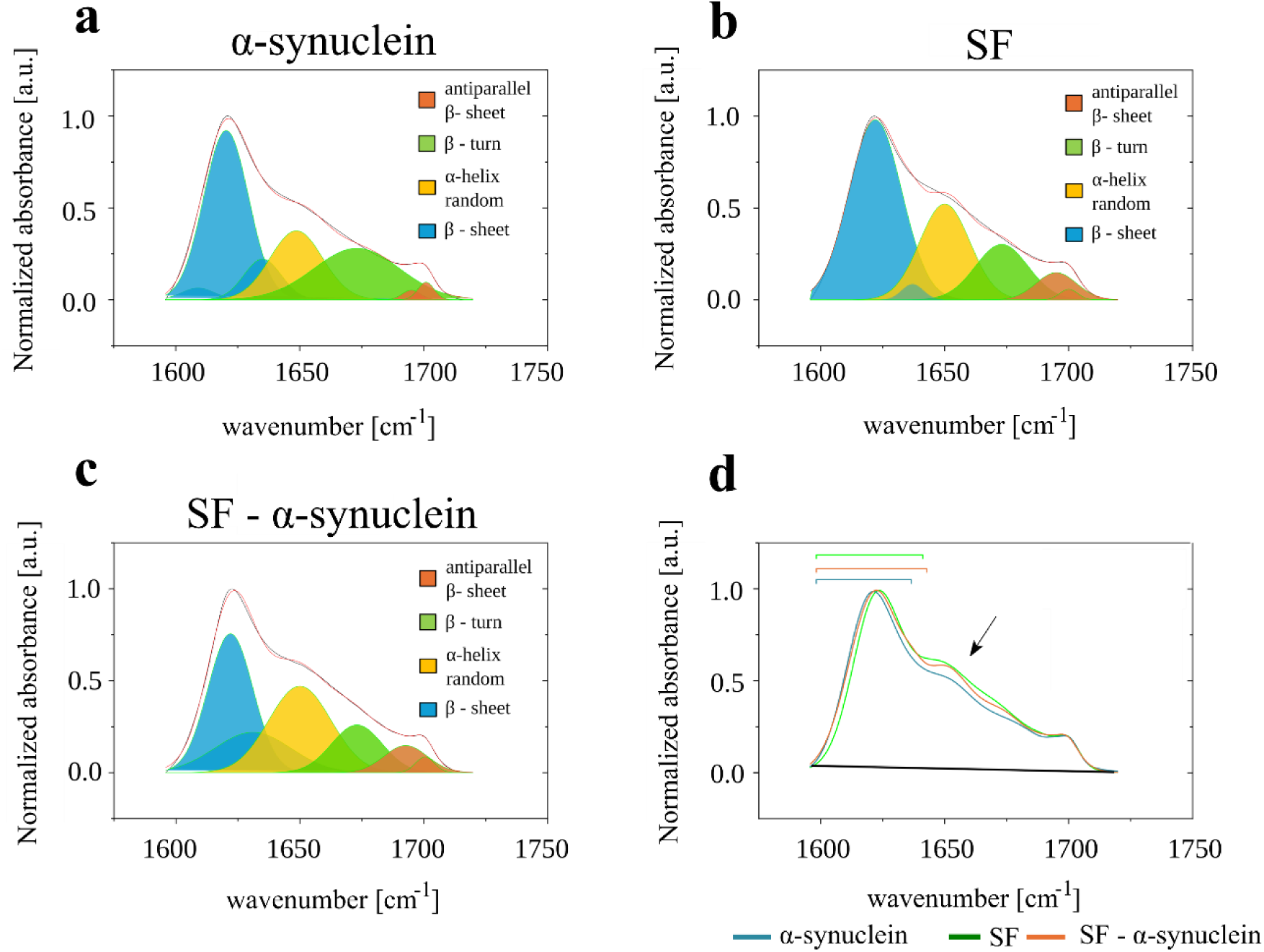
Secondary structure analysis by Fourier transform infra-red (FTIR) spectroscopy. FTIR analysis has been performed for **a)** α-synuclein, **b)** SF and **c)** SF- α-synuclein assemblies. The quantitative assessment of the secondary structures was assigned as follow: 1609, 1621, and 1631 cm^−1^ for intermolecular β-sheets (shown in blue); 1650 cm^−1^ for α-helix and random coil (shown in yellow), 1673 and 1703 cm^−1^ for β-turn (shown in green); and 1695 cm^−1^ for antiparallel amyloid β-sheets (shown in red) (**d**) Graph showing and overlap between FTIR spectra collected for amide I region of α-synuclein, SF and SF- α-synuclein assemblies.

**Supplementary Figure 9.**
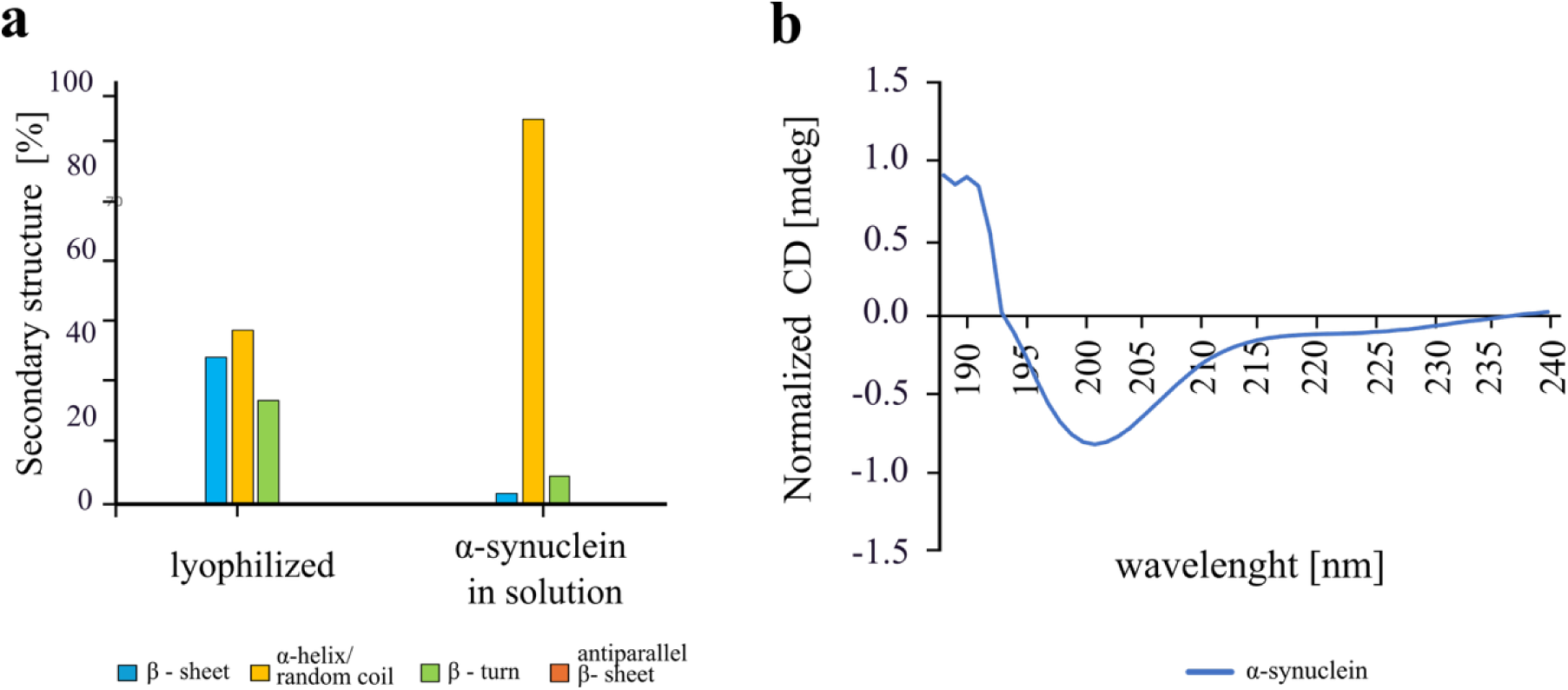
Secondary structure analysis of the soluble and aggregated α-synuclein structures. **a)** Chart summarizing secondary structure content of α-synuclein in it’s different solubility states (in solution and lyophilized) by using Circular Dichroism (CD) and FTIR spectroscopy. **b)** Graph showing the original CD spectrum of α-synuclein protein in buffer (see Methods).

